# Enhanced capillary pumping using open-channel capillary trees with integrated paper pads

**DOI:** 10.1101/2022.07.15.500252

**Authors:** Jodie C. Tokihiro, Wan-chen Tu, Jean Berthier, Jing J. Lee, Ashley M. Dostie, Jian Wei Khor, Madeleine Eakman, Ashleigh B. Theberge, Erwin Berthier

**Affiliations:** Department of Chemistry, University of Washington, Seattle, Washington 98195, USA; Department of Urology, University of Washington, Seattle, Washington 98195, USA

**Author notes:** J.C.T., W-c. T., and J.B. contributed equally to this work.

## Abstract

The search for efficient capillary pumping has led to two main directions for investigation: first, assembly of capillary channels to provide high capillary pressures, and second, imbibition in absorbing fibers or paper pads. In the case of open microfluidics (i.e., channels where the top boundary of the fluid is in contact with air instead of a solid wall), the coupling between capillary channels and paper pads unites the two approaches and provides enhanced capillary pumping. In this work, we investigate the coupling of capillary trees— networks of channels mimicking the branches of a tree—with paper pads placed at the extremities of the channels, mimicking the small capillary networks of leaves. It is shown that high velocities and flow rates (7 mm/s or 13.1 µL/s) for more than 30 seconds using 50% (v/v) isopropyl alcohol, which has a 3-fold increase in viscosity in comparison to water; 6.5 mm/s or 12.1 µL/s for more than 55 seconds with pentanol, which has an 3.75-fold increase in viscosity in comparison to water; >3.5 mm/s or 6.5 µL/s for more than 150 seconds with nonanol, which has an 11-fold increase in viscosity in comparison to water) can be reached in the root channel, enabling higher sustained flow rates than that of capillary trees alone.

## I. INTRODUCTION

Simple and autonomous microfluidic systems can be designed by use of capillary forces. In such systems, bulky active pumps^1,2^ are not needed. However, a major drawback in a capillarity-based system is the decrease of the flow velocity—and the flow rate—with time. According to the Lucas-Washburn-Rideal (LWR) law, the decrease of the velocity is proportional to the inverse of the square root of time.^3–5^ To overcome this drawback, different capillary pump designs have been developed.

Many passive designs have been developed for capillary-driven flows in closed channels ^6–9^; however, few have been proposed in the case of open channels. The electrowetting-based pumping device proposed by Satoh *et al.* is one of the first pumping designs in open geometries, but requires the addition of electric actuation.^10^ The most current open-pumping systems rely on evaporation. Evaporation from a rectangular open channel has been documented by Kolliopoulos *et al.*^11^ and Lynn *et al.*^12^ and pumping has been set from a reservoir or fibrous pads by Zimmermann *et al*.^13^ However, these methodologies are restricted to low boiling point liquids and entail long experiment durations. The search for efficient open channel pumping based solely on geometrical features is currently progressing. Srinivasan has proposed a geometrical diffuser for zero gravity pumping in space vanes^14^ and Guo *et al.* have developed an interesting system combining capillarity in closed channels and additional pumping from paper pads has been documented.^15^

In this work, the pumping mechanism involves the use of networks of small open channels where the capillary pressure is high, or there are matrices of fibers (often paper pads) with a high wicking power. These capillary pumps are placed behind the “region of interest” where the biological or chemical processes are performed and can be used in multiple applications including separation methods or sample processing.

Capillary trees for capillary pumping in closed (confined) channels have been developed in the past.^16^ Examples include triple tree line capillary pumps used for performing immunoassays,^17^ microstructures for simple and advanced capillary pumping,^18^ and multilayers of microfluidic paper to generate the capillary flow.^19, 20^ On the other hand, it was shown that microporous and fibrous structures, such as paper pads or porous membranes, provide efficient pumping properties due to their high wicking power.^21–23^

In the capillary-driven microfluidics field, open systems are of special interest.^24–26^ These systems remove at least one “wall” of the microfluidic channel (often the top wall), providing easy access to the flowing liquid in the channels. We have previously found that a capillary tree can be used to maintain a high value of velocity in the root channel—the channel of interest for a given application—in open microfluidic devices.^27^

In this work, we show that these open capillary tree channels can easily be connected to paper pads (mimicking the small capillary networks of leaves) to further extend capillary pumping in the root channel which is an innovation from our previous work^28^. Extending periods of high flow velocity is an important problem in open capillary microfluidics. Here, we have chosen to use an open channel instead of a closed capillary tree due to fabrication limitations such as difficulty in adjusting a cover plate over the trees with minimal to no leakage of the flowing solvent. Along with our previous work, this work expands the toolbox for open-channel microfluidics, which is implemented in a broad range of fields of studies such as diagnostics, biomedical science, pharmaceutical science, space science, and integrated analysis systems. The removal of the top wall of the channel enables direct access to the flowing fluid for the addition, removal, or manipulation of the channel contents. Thus, together with this work and prior work,^28^ we demonstrate a method for sustaining periods of high fluid velocity through the coupling of paper pads (current work) and bifurcating capillary trees (prior work). In itself, an “infinite” capillary tree with decreasing channels has been shown to be favorable for pumping, but practically, it is impossible to design and fabricate such a device. In addition, paper alone is limited by the high friction of the liquid in the microporous media. Hence, the additive structure constituted by the paper pads replaces the downscaling of the trees and enhances the global pumping effect, first coming from the capillary tree and then from the fibrous paper “leaves”. The paper pads are placed in milled receptacles at the end of the branched channels. We used a geometric design of capillary trees where the root channel is successively divided in a cascade of daughter channels and the cross sections of the daughter channels are progressively decreased at a ratio of 0.85. With this design, a high flow rate in the root channel can be achieved and sustained with such designs using bifurcations and paper pads (Fig. 1). The multiplication of the tree branches avoids the velocity decrease in the root channel predicted by the LWR law in the case of a constant cross-section channel. The flow rate decreases in the branches after each bifurcation, but the flow rate in the root channel is twice that of the first branches and four times that in the second branches, and so on. On the other hand, the multiplication of the pads in parallel palliates the friction effect in each pad on the flow rate in the root channel.

**FIG. 1.**
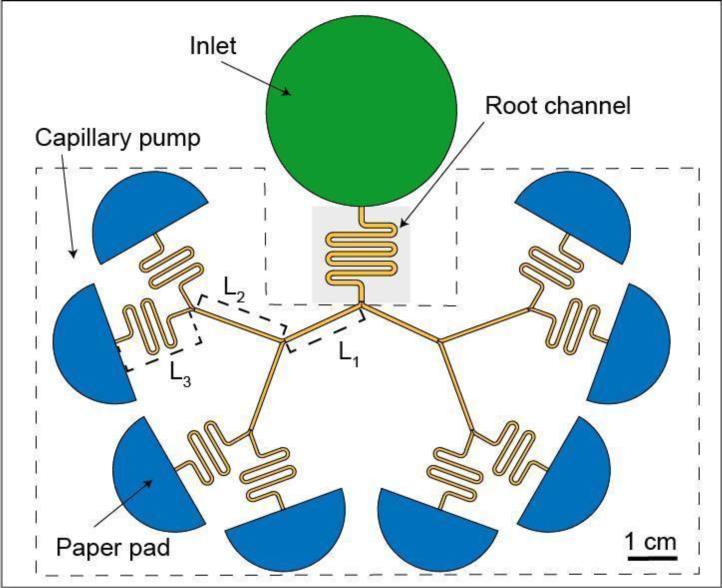
Diagram of an open channel microfluidic device utilizing a capillary tree channel and paper pads.

A closed form model for the flow dynamics is derived here, coupling the formulation of the capillary tree flow with the paper pads. In this manuscript, it will be shown that high capillary velocities are obtained even in the case of highly viscous fluids, pentanol, and nonanol. While there are publications where paper pads are used to drive flow^15, 23^, to the best of our knowledge, this is the first study to couple paper pads to bifurcating trees, specifically in open channel systems. We show here that by combining homothetic capillary tree channels with paper pads in an open microfluidic device, such designs maintain a high liquid velocity (7 mm/s or 13.1 µL/s) for more than 30 seconds using 50% (v/v) isopropyl alcohol, which has a 3-fold increase in viscosity in comparison to water; 6.5 mm/s or 12.1 µL/s for more than 55 seconds with pentanol, which has an 3.75-fold increase in viscosity in comparison to water; >3.5 mm/s or 6.5 µL/s for more than 150 seconds a fluid that has a viscosity that is 11 times higher than water in the root channel for more than one minute.

## II. MATERIALS AND METHODS

### 2.1. Fabrication of capillary tree channels

The device consists of winding serpentine-shaped channels (the root channel), a large inlet in which the liquid is introduced using a pipette, three levels of branches, and semicircle paper pads at the extremity of the last set of channel branches (Fig. 1). The dimensions of the channels are listed in Table SII; an engineering drawing is included in Figure SI; and the computer-aided design files are included in the electronic Supplementary Material. The widths and depths of the channels are homothetically reduced by a factor of 0.85 after each bifurcation. The turns in the winding channels do not affect the capillary flow in the absence of capillary filaments,^29^ as the rounded bottom avoids the formation of filaments observed in channels of rectangular cross section.^30^ The average wall friction length of the root channel is estimated to be λ∼ 259 µm from our preceding work.^27, 28^ It was shown that the average friction length produces the value of the average wall friction, τ, by the formula, τ = μ *V*/λ^21^, where *μ* is the liquid viscosity.

The device was designed using a computer aided design (CAD) software (Solidworks 2017, Waltham, MA) and the design files were converted to G-code using a computer aided manufacturing (CAM) software (Fusion 360). Channels were milled in poly(methyl methacrylate) (PMMA) sheets (3.175 mm thick, #8560K239; McMaster-Carr, Santa Fe Springs, CA). To create round bottom channels, endmills with a cutter diameter of 1/32” (TR-2-0312-BN) and 1/64” (TR-2-0150-BN) were used (Performance Micro Tool, Janesville, WI).

The devices were fabricated via micromilling on a Datron Neo computer numerical control (CNC) mill (Datron, Germany). The channel bottom is estimated to have a few microns of roughness which is one magnitude below the roughness values that were observed by Lade et al.^31^ that would produce fluctuations in flow velocity.

### 2.2. Paper pads

Whatman #1 paper (Whatman Grade 1 Qualitative Filter Paper, #28450-160, VWR Scientific, San Francisco, CA) was cut into half circle shapes using a plotter cutter (Graphtec Corporation, Yokohama, Japan). A tight contact between the paper pads and the outlets of the last set of tree channels is essential to maintaining the capillary flow of the fluid. The main characteristics of the paper pads are listed in the Supplementary Material Table SIII. The capillary pressure depends on the liquid that is used and the values are in alignment with the experimental results.

### 2.3. Solvents

The physical properties of the solvents are indicated in the Supplementary Material Table SI. To mitigate evaporation of the solvents, pentanol and nonanol, which are low volatile solvents, (boiling points are 139°C and 213°C for pentanol and nonanol, respectively), were used. Both pentanol and nonanol have been colored with either Solvent Yellow 7 or with Solvent Green 3 (Sigma-Aldrich) at concentrations of 0.50 mg/mL and 1.43 mg/mL, respectively. Aqueous isopropyl alcohol (IPA) (VWR Scientific) was used at a concentration 50% (v/v) and colored with 0.60% yellow or 1.2% blue food coloring. (McCormick).

### 2.4. Capillary trees with integrated paper pads flow experiments

To obtain fluid travel distance and velocity data, 2 mL of the yellow-dyed fluid was pipetted into the reservoir of the device. Once the fluid front reached the end of level 1, a 200 µL refill of the yellow fluid was added. After the yellow fluid wetted the paper pad, 500 µL of the blue-dyed fluid was pipetted into the fluid reservoir with a 200 µL refill after the blue fluid front reached the end of level 1. Data was reported up to the point with when the paper pad was saturated at 90 s for 50% (v/v) isopropyl alcohol, 57 s for pentanol, 154 s for nonanol, for trials 1-3, respectively (Fig 3 and Fig. SII-SIII).

### 2.5. Imaging and Analysis

Videos of the progression of the solvent flow in the device were recorded using a Nikon-D5300 ultra-high resolution single lens reflective (SLR) camera. The images of the videos were extracted automatically by the code. The location of the tip of the 50% IPA flow was pinpointed using MATLAB software, while the travel distance of the pentanol and nonanol flows were measured manually using Fiji (ImageJ) software. For the manual analysis, the scale was set for an individual trial with the “Set Scale” tool. The fluid front was pinpointed with the “Segmented Line” tool and the “Measure” function was used to calculate the total distance traveled along the capillary tree. Each data was from every 10 frames (yellow) or 30 frames (blue).

## III. RESULTS AND DISCUSSION

### 3.1. Theory

In the first phase, the flow advances in the capillary tree, dividing itself at each bifurcation. The device is designed so that the tree is symmetrical, thus all fluidic channels have the same length. This motion has already been analyzed in our prior work.^27^ We describe here a theoretical approach for open capillary channel bifurcating trees coupled to paper pads. The nomenclature and associated definitions used in this section can be found in Table I.

**TABLE I.** Nomenclature for theoretical approach.

| Symbol | Definition | [Unit] |
| --- | --- | --- |
| Latin, lower case |  |  |
| $p$ | perimeter | [mm] |
| $p_n$ | perimeter of root channel | [mm] |
| $p_n$ | perimeter of branch n | [mm] |
| $t$ | time | [s] |
| $t_n$ | time at extremity of branch n | [s] |
| $t_0$ | time at exit of the root channel | [s] |
| $z$ | travel distance | [mm] |
| $z_0$ | travel distance (root channel) | [mm] |
| $z_n$ | travel distance in branch n | [mm] |
| $z_p$ | travel distance in the paper pad | [mm] |
| Latin, upper case |  |  |
| $A_n$ | characteristic treelength (n branches) | [mm] |
| $C$ | coefficient of the LWR law | [mm<sup>2</sup>/s] |
| $K$ | permeability (paper pad) | [mm<sup>2</sup>] |
| $L$ | length of a branch | [mm] |
| $L_0$ | length of root channel | [mm] |
| $P$ | pressure | [Pa] |
| $P_j$ | pressure at node j | [Pa] |
| $P_{cap}$ | capillary pressure | [Pa] |
| $S$ | cross-sectional area | [mm<sup>2</sup>] |
| $S_0$ | cross-sectional area (root channel) | [mm<sup>2</sup>] |
| $S_{p0}$ | cross-sectional area (paper pad entrance) | [mm<sup>2</sup>] |
| $V$ | velocity | [mm/s] |
| $V_0$ | velocity (root channel) | [mm/s] |
| $V_n$ | velocity (branch n) | [mm/s] |
| Greek, lower case |  |  |
| $\alpha$ | homothetic factor | [non-dimensional] |
| $\beta$ | geometrical angle of the circular pad | [rd] |
| $\gamma$ | liquid-air surface tension | [mN/m] |
| $\phi$ | porosity (paper pad) | [non-dimensional] |
| $\lambda$ | friction length | [mm] |
| $\mu$ | liquid viscosity | [mPa.s] |
| $\tau$ | time (for paper pads) | [s] |
| $\theta$ | Young angle | [rd] |
| $\theta^*$ | generalized Cassie angle | [rd] |
| Greek, upper case |  |  |
| $\nabla$ | gradient operator | [1/mm] |

Let us recall that the marching distance in the open root channel (the channel before the beginning of the bifurcations) is given by

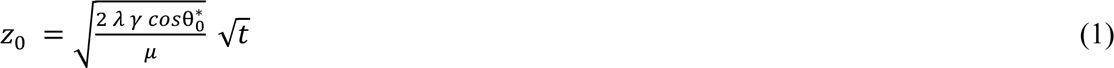

where λ is the average wall friction length,^29^ γ is the surface tension, *μ* is the viscosity, *t* is the time, *θ\** is the generalized Cassie angle,^32^ and index, *0*, refers to the root channel, so that 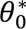 is the generalized Cassie angle in the root channel. Using the pressure at each node (bifurcation) plus the homothetic relation for the channel perimeters, and cross sections and the mass conservation equation, one finds the expression of *z_n_* for the marching distance in the channels after the n^th^ bifurcations:

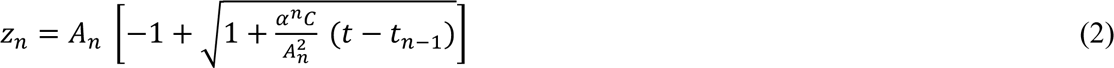

where 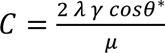, α is the homothetic ratio, t_n-1_ is the time at which the liquid enters the nth channel, and *A_n_* is a geometrical factor which depends on the channel lengths *L_0_* to *L_n-1_* and α. The algebra leading to this expression is lengthy and fully developed in the Supplementary Material Section 1. Note that (2) differs from the LWR expression where *z* ∼ *t*^1^*^/^*^2^.

Fig. 2 (Multimedia View) shows the flow of fluid in a device that combines a capillary tree and paper pads. When the flow reaches the paper pads, the wicking of the pads is governed by Darcy’s law.^33, 34^

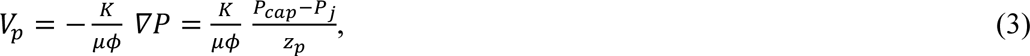

where *P_cap_* is the capillary pressure of the paper, *P_j_* is the pressure at the channel-paper pad junction, *K* is the permeability of the pad, and *φ* is its porosity. The index, *p,* refers to the paper, and the triplet (*P_cap_*, *K*, *φ*) characterizes the paper strip.^35, 36^ The capillary pressure of the paper pad must be larger—in absolute value—than the pressure, *P_j_*, at the last nodes so that the liquid continues to flow in the pads (*V_p_* >0). This condition is easily satisfied even if the capillary pressure in the tree increases after each bifurcation: 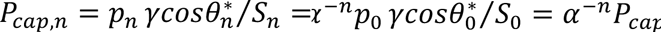. If we remark that *α*^-*n*^*P*_*cap*,0_∼50 Pa, and that the capillary pressure in most paper pads is of the order of 3000 Pa at least (Supplementary Material Table SIII), more than 25 bifurcations will still produce a capillary pressure inferior to that of the pad—considering *α* = 0.85. The derivation of the flow motion in the pads (coupled to the tree) is detailed in the Supplementary Material Section 2. Note that three assumptions are used in the model. First, that the paper pads are homogeneous (i.e., there is no region of higher or lower porosity). Hence, saturation is neglected, and the sharp front assumption is used.^37–39^ This observation is confirmed by experiments.^23, 37^

**FIG. 2.**
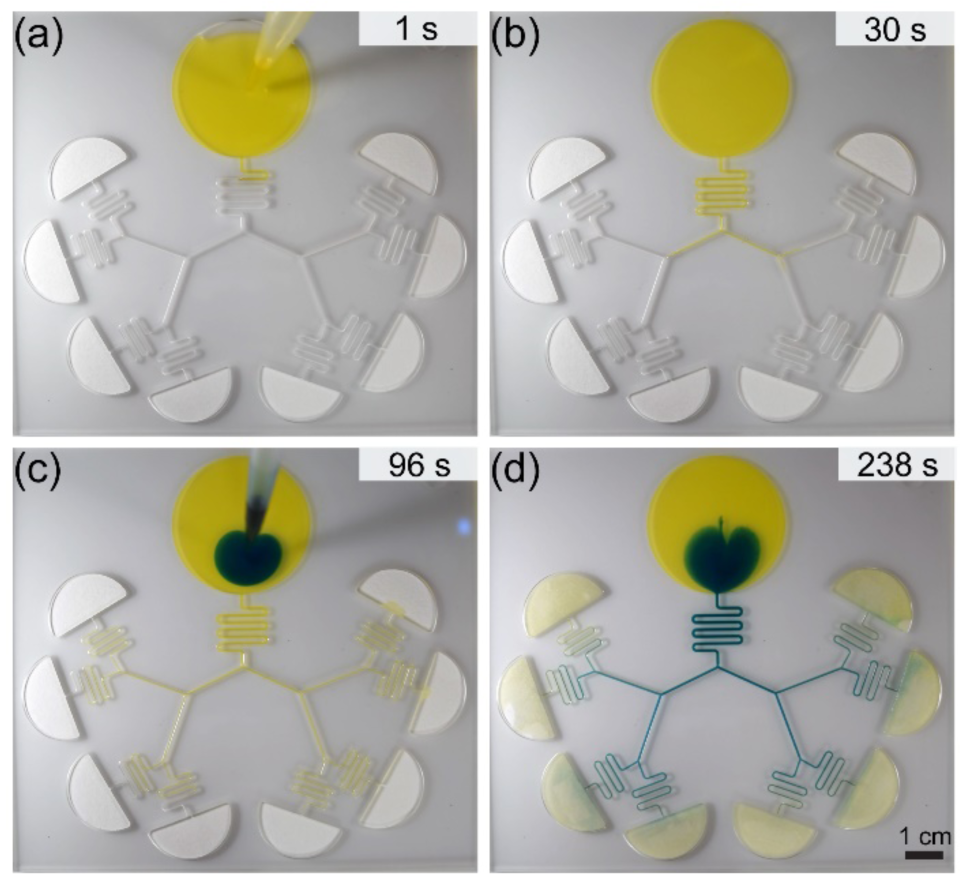
Still images of an open channel device filled with yellow and blue nonanol solutions. (a) Yellow liquid is pipetted in the inlet of the open channel. (b) Progression of the yellow liquid in the root channel and capillary tree. (c) Blue liquid is pipetted in the inlet after the yellow liquid reaches the paper pad. (d) Progression of blue liquid in the open channel. Scale bar is 1 cm. (Multimedia View).

Second, it is assumed that the dilatation of the paper fibers from the liquid is negligible, therefore, the porosity, *φ,* is constant everywhere in the pad. Third, the cellulose fibers do not absorb the wicking liquid, thus the mass conservation of the flowing liquid is independent of time.

Equation (3) can be solved using the expression of the pressure, *P_j_*, found in the first phase (flow in the tree) and assuming a circular or flat contact line in the conical (angle *β*) or rectangular pads (*β* = 0). The travel distance in the paper is then

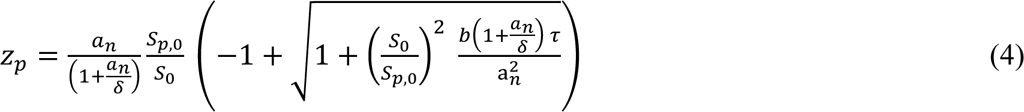

where 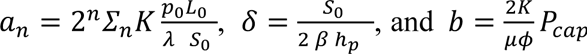. Here, *p_0_* corresponds to the total perimeter of the cross section of the root channel, *S_0_* is the cross-section surface area, *h_p_* is the thickness of the paper pad, *β* is the paper pad cone angle, *τ* is the time counted from the moment when the liquid reaches the pads, and 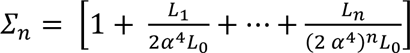. Note that if *β* is zero, then *δ* is infinite, resulting in 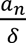 to cancel out in equation (4). The two first parameters, *a*_*n*_ and *δ*, have the dimension of length, while the unit for *b* is mm^2^/s and *Ʃ_n_* is dimensionless. The ratio, 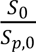, is the ratio between the cross-sectional area of the root channel and that of the paper pad (at the junction with the tree). Using the mass conservation equation, the velocity in the root channel when the liquid wicks the paper is given by

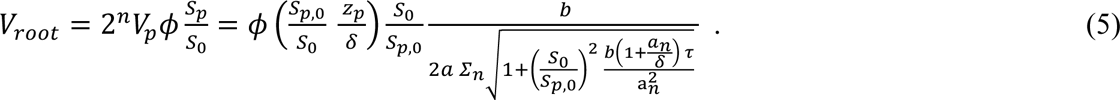

where *z_p_* is given by (4).

### 3.2. Comparison with experiments

The travel distance produced by equation (S13) (Supplementary Material Section 2) has been checked against the experiments using colored solutions of 50% IPA (v/v), pentanol, nonanol (Fig. 3) flowing in the homothetic tree of ratio *α* = 0.85. A representative trial for each fluid is shown in Fig. 3. Raw data for the travel distance for each fluid is presented in Supplementary Material Fig. SII. The travel distances in the tree, as measured by the progression of the fluid front, are well matched by the theory. The theory assumes a perfect flow with a circular fluid front in the semi-circular paper pad and perfect synchronization between the pads; however, in our experiments, the fluid front deviates from the theory due to a not-perfectly circular liquid front in the paper pad, imperfect synchronization between the flows in the different branches, and occasional leaking in the space below the pads. Nonetheless, this does not preclude the data. Note that a velocity jump is predicted by the model at the tree/pad junction due to the sudden change of capillary pressure. However, this velocity jump is not high due to the friction along the whole capillary tree. For this reason and because of the connection between channel and pads, this jump is hardly seen in the experiments.

**FIG. 3.**
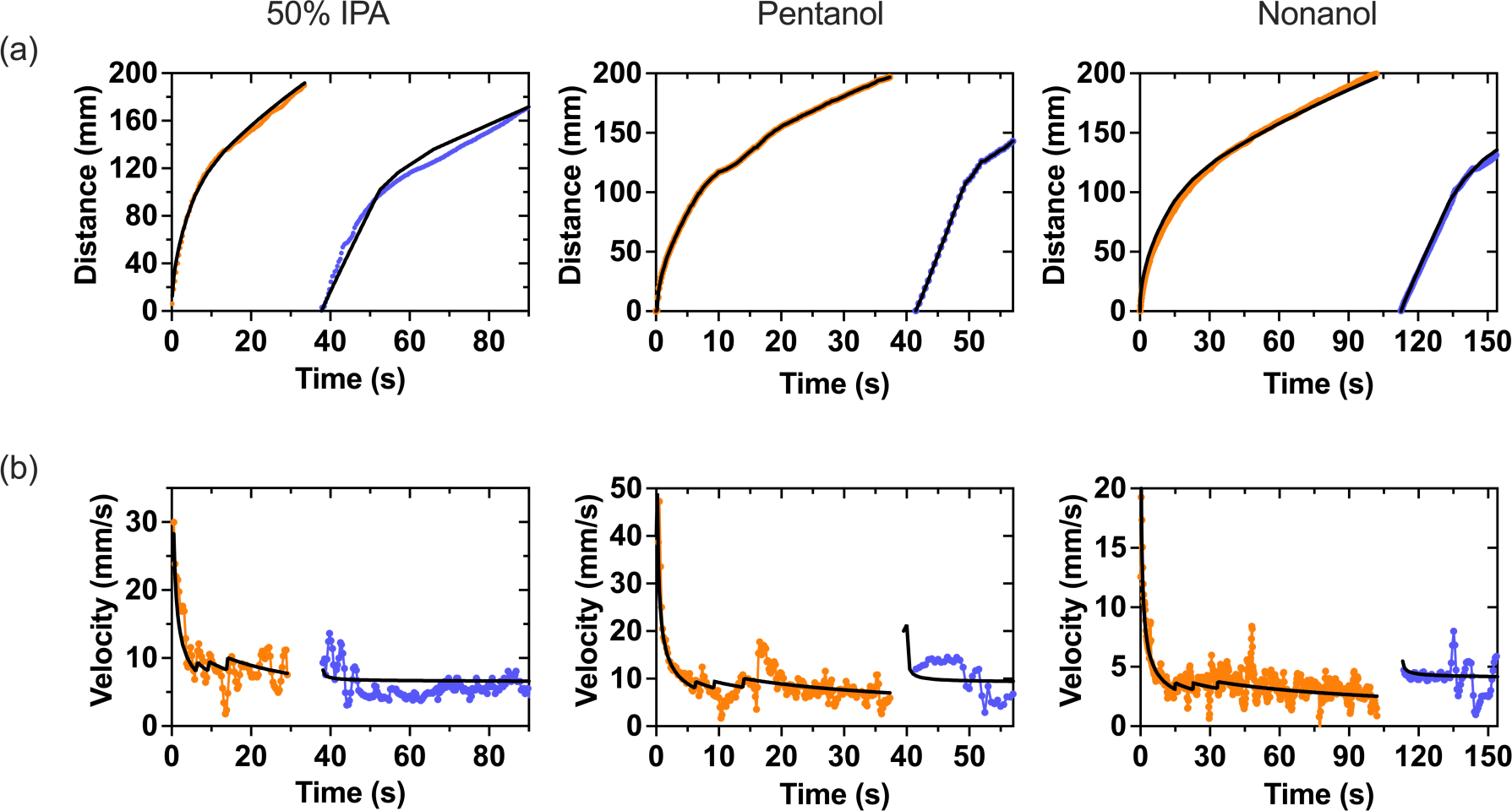
Travel distance and velocity over time for 50% (v/v) IPA solution, pentanol, nonanol solutions in an open channel capillary tree with paper pads. (a) Experimentally determined travel distances in the bifurcating capillary tree vs. time for the first phase of the flow (orange dots) and the second phase of the flow (blue dots) after the flow has reached the pads and blue colored solution has been added to the inlet. Travel distance was determined by measuring the distance of the fluid front in the device in the recorded videos. The black lines are the theoretical results. (b) Velocities in the root channel vs. time when the tip of the flow is in the capillary tree (orange dots) and in the paper pads (blue dots). Velocities were determined using equation (5).

In the case of 50% IPA, the coefficient, 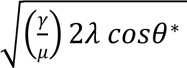, was found to be approximately 40 mm/s^1/2^ (Supplementary Material Table SI). In the paper pads, a good fit was found for a capillary pressure of 3000 Pa. Root channel velocities on the order of 7 mm/s were obtained for 60 seconds (Fig. 3) for 50% IPA. The contact of the liquid with the pads occurs at 35 seconds for both 50% IPA and pentanol, and 95 seconds for nonanol (Fig. 3). Raw velocity data plots are presented in Supplementary Material SIII.

For the case of pentanol, which is a fluid with an “intermediary” viscosity of 3.75 mPa.s. the pentanol also has a good fit for the travel distances in the capillary tree is obtained for the value 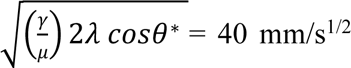. Root channel velocities on the order of 6.5 mm/s were obtained for at least 55 seconds (Fig. 3).

The case of nonanol is of great interest due to its high viscosity of 0.011 Pa.s. A very good fit for the travel distances in the capillary tree is obtained for the value 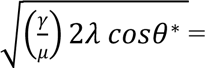 23.7 mm/s^1/2^, and a capillary pressure *P_cap_* = 5500 Pa in the paper pad. Root channel velocities on the order of 3.5 mm/s were obtained for 150 seconds (Fig. 3).

The fluctuations that were observed in the velocity measurements (Fig. 3) are due to the discretization, *d*z/*d*t. The velocity is the derivative of the travel distance and derivation amplifies the fluctuations. The only physical variation is at the junction between the channel and the paper due to abrupt capillary pressure changes.

### 3.3 Discussion

Efficient capillary pumping has been the subject of many investigations. In this problem, three parameters must be considered: (1) the maximum velocity of the flow, (2) the maximum flow rate, and (3) the duration of the pumping. Prior research has focused on obtaining the highest possible velocities, but typically only for a short time and a moderate volumetric flow rate. For example, Reches *et al.* ^40^ has obtained interesting velocities of 2 cm/s for water in treated threads of wool, but along a length of 2 cm, corresponding to a duration time of 1 second. In Table II, a review of the literature for various methods of capillary pumping is summarized. The table indicates that it is difficult to obtain high velocities (larger than 1 mm/s) for a long duration (larger than 30 seconds) in capillary-based systems.

**TABLE II.**
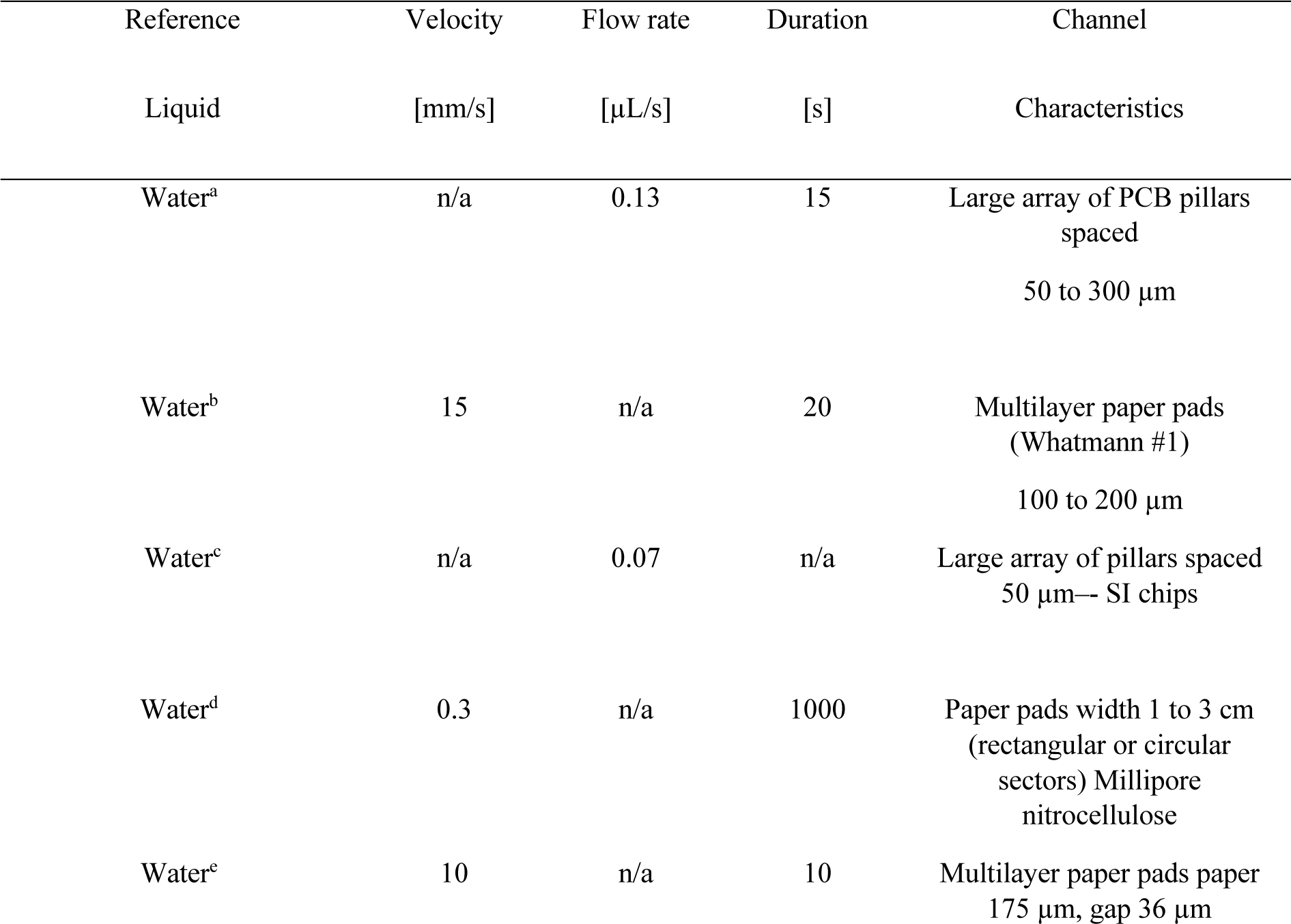
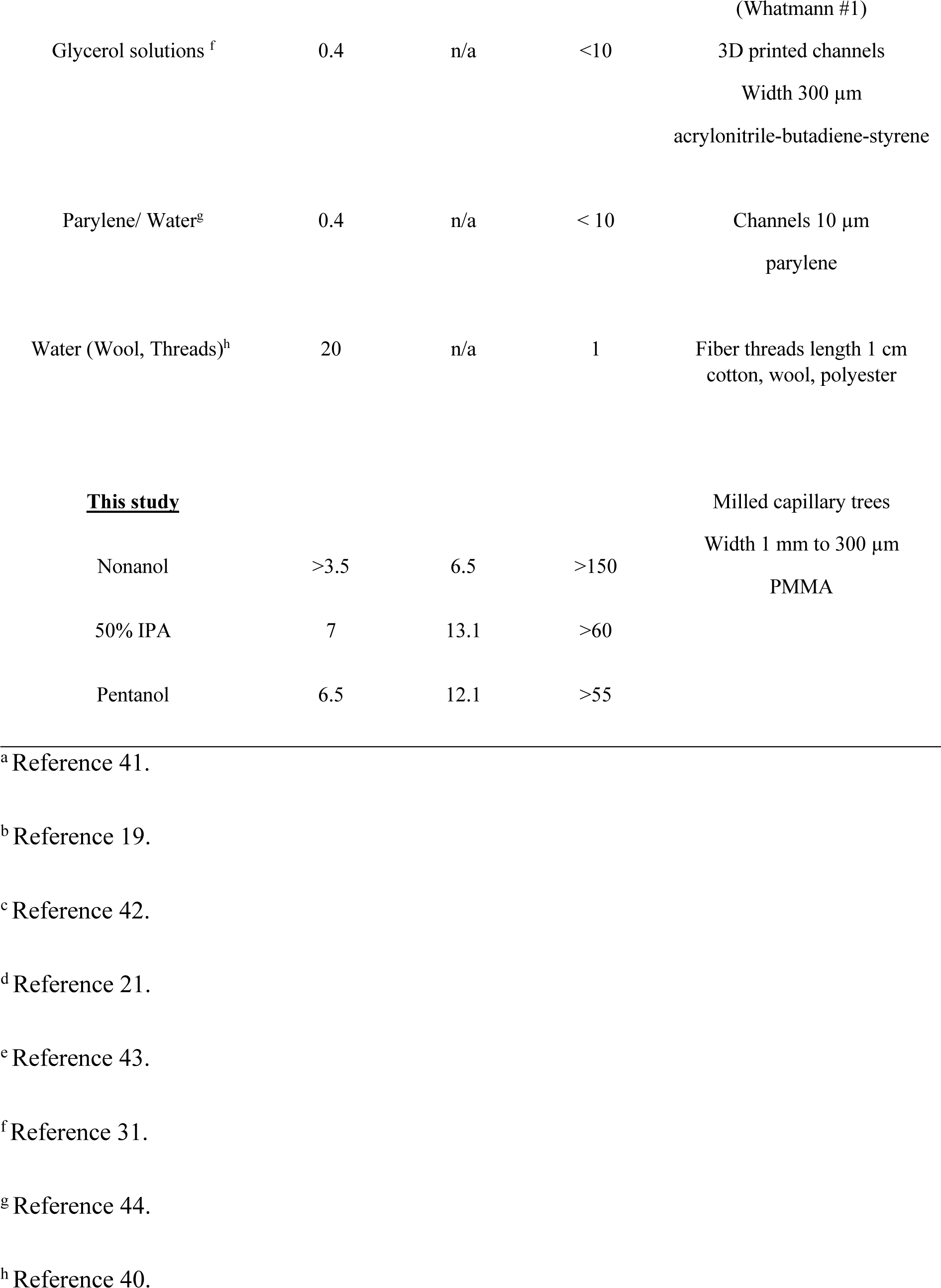
Literature review of capillary pumping.

In our study, it is assumed that the capillary tree is “symmetrical”, i.e., all branches at the same level of ramification are identical. The channels are relatively larger in size due to CNC milling constrictions; however, this theory is applicable to smaller dimensions up to 300 µm in width when employing other types of fabrication methods. High velocities are obtained with our device where the dimensions are at the upper side of the microscale limits. If we remark that the root channel velocity is approximately proportional to the square root of the friction length and that the friction length decreases proportionally with the channel dimension, a homothetical reduction of the channel dimension of a factor, n, will result in the reduction of the velocity of a factor 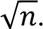. For example, if the channel cross-section is decreased by 4 (200 µm width), the velocity will be reduced by a factor of 2. Still, high velocities are obtained with the device for microscale channels. The progression of the liquid is the same for each path and is obtained by using the extended LWR law that states that the dynamics of the flow results from the balance between the capillary force on the advancing meniscus and the wall friction along the path.^3–5, 24, 25^ Intrinsically, capillary-driven flow velocities decrease as capillary length decreases. Here, our device enables longer capillary lengths in the root channel due to the multiplication of daughter branches while maintaining high flow velocities, allowing for applications in diagnostics and bioanalytical chemistry.^24^ Note that the approach proposed here is also applicable to closed systems by using the friction length of the closed channels instead of open capillary channels.^24, 25^ In the microporous paper pads, the wicking liquid velocity is given by Darcy’s law^33, 34^ and determined by three parameters: permeability, porosity, and capillary pressure.^35, 36^

To validate the closed form model derived in this work, experiments were performed using nonanol, isopropyl alcohol, and pentanol solutions in open microfluidic channels. We chose these liquids because they wet native PMMA without surface treatment (θ = 13 - 47°, Supplementary Material Table SI). Further, evaporation of these liquids is slow compared to the time scale of the capillary flow. Hence, evaporation is not taken into account for this study.^3, 45–47^ High capillary velocities are obtained even in the case of highly viscous nonanol, which has a viscosity eleven times that of water (Supplementary Material Table SI). In the case of the less viscous IPA aqueous solution, velocities higher than 1 cm/s were obtained.

The device presented here is of interest for its ability to combine velocity, flow rate, and duration. This innovation builds on our our previous work^28^ to further enable extended periods of high velocity flow in open microfluidic channels, which is desired for capillary microfluidics research and device development for fields such as chemical, biomedical, biological, space science, and materials research. In the future, the device can be optimized to achieve higher velocity. A longer length of the last branch of the capillary tree can enable a higher velocity in the root channel. The choice of the paper matrix is of great importance, especially the two parameters *K/φ* (Leverett parameter) and capillary pressure (*P_cap_*). Longer pads would allow longer duration of the high velocity flow. Additionally, the device can be micromilled using different materials, such as polystyrene, which enables wider application.^48^ Devices can be oxygen plasma treated to allow flow of aqueous solutions (cell culture media, biological fluids, etc.).^26, 48^

## IV. CONCLUSION

In this work, the dynamics of the capillary flow circulating in a capillary tree with paper pads placed at the extremities of the capillary tree branches have been investigated. A model for the dynamics of the flow in the capillary tree has been coupled to a model for the flow in the paper pads. This coupling has been validated against experiments performed with milled open PMMA channels and paper pads. It is first shown that capillary trees with homothetically decreasing cross-sectional areas (in a ratio of 0.85) maintain the flow velocity in the root channel. Moreover, the presence of paper pads at the extremities of the branches prolongs the duration of the high flow rate pumping. The present analysis demonstrates the possibility of obtaining high velocities and flow rates (7 mm/s or 13.1 µL/s) for more than 30 seconds for 50% IPA, 6.5 mm/s or 12.1 µL/s for more than 55 seconds with pentanol, and flow rates of >3.5 mm/s or 6.5 µL/s for more than 150 seconds for nonanol. These flow rates are nearly constant (save for periodic jumps due to experimental fluctuations) if conical-shaped paper pads are used as suggested in the literature.^49, 50^ For volatile fluids, the employment of evaporation could extend the duration of these high flow rates. Further, we envision many areas of future application including using our method to push the limits of viscous fluid flow in open channels as we can enable the sustained passive flow of complex biological fluids such as saliva or blood. This device also has the potential to be applied to biological experiments such as *in vitro* cell culture or analytical methods involving biological fluids. Further, the method presented here can be adapted for diverse applications of open microfluidics including chemical synthesis, chemical analysis, thermics, material science, or space science, allowing for diverse applications at the frontiers of science.

## SUPPLEMENTARY MATERIAL

See supplementary material for:

- Open capillary padding design file
- Additional information on physical properties of solvents and paper pads, channel dimensions, theoretical derivations, and engineering drawing of microfluidic device.

## ACKNOWLEDGEMENT

This work was funded by NIH grant R35GM128648. The content is solely the responsibility of the authors and does not necessarily represent the official views of the National Institutes of Health. We also acknowledge the M.J. Murdock Foundry for Translational Research. We would also like to thank David N. Phan for writing the code for video extraction.

