## Supplementary Material for "Enhanced capillary pumping using open-channel capillary trees with integrated paper pads"

**Additional supporting information provided**

- Design file: Open capillary padding (.SLDPRT).

TABLE SI. Physical properties of the solvents. The term $\sqrt{\left(\frac{\gamma}{\mu}\right)2\lambda cos\theta^*}$ is the coefficient of the extended Lucas-Washburn law and there is a slight difference between the value obtained from literature and from experiments.

| Physical property | 50% IPA | Nonanol | Pentanol |
|---|---|---|---|
| [a] Surface tension $\gamma$ [mN/m] | 24 | 28.5 | 25.4 |
| [a] Viscosity $\mu$ [mPa•s] | 3.2 | 11.2 | 3.75 |
| [a,b] Contact angle $\theta$ [deg] | 18 | 13 | 10 |
| $\gamma \cos \theta$ [mN/m] | 22.8 | 27.8 | 25.5 |
| [c] $\gamma \cos \theta^*$ [mN/m] | 14.6 | 15.3 | 15.5 |

| | | | |
|---|---|---|---|
| [d] Literature data $\sqrt{\left(\frac{\gamma}{\mu}\right) 2\lambda\, cos\theta^*}$ [mm/s$^{1/2}$] | 48.2 | 28.5 | 38.7 |
| [e] Experimental data $\sqrt{\left(\frac{\gamma}{\mu}\right) 2\lambda cos\theta^*}$ [mm/s$^{1/2}$] | 40 | 23-25 | 40 |

[a] air at room temperature[4-7]; [b] contact angle of the solvent on native PMMA; [c] $cos\ \theta^*$ is the Cassie contact angle[8]; [d] calculated using values in the table on line 1 ($\gamma$), 2 ($\mu$), 5 ($\gamma\ cos\ \theta^*$), and $\lambda$ = 259 µm[1,2].

[e] fit with the travel distance in the root channel ($z_0$).

TABLE SII. Characteristic dimensions of the homothetic channels

| Dimension | Level 0 (root) | Level 1 | Level 2 | Level 3 |
|---|---|---|---|---|
| Width [mm] | 1.06 | 0.90 | 0.77 | 0.65 |
| Depth [mm] | 1.76 | 1.50 | 1.28 | 1.08 |
| Wetted perimeter $p_0$ [mm] | 4.13 | 3.51 | 3.00 | 2.53 |
| [a]Friction length $\lambda$ [µm] | 259 | 220 | 187 | 159 |
| Cross-sectional area $S_0$ [mm$^2$] | 1.75 | 1.26 | 0.92 | 0.66 |

[a] obtained from previous publication.[1,2]

TABLE SIII. Characteristics of Whatman #1 paper pads.

| [a] Physical and geometrical properties | Symbol | Value | Unit |
|---|---|---|---|
| Permeability | $K$ | 2.3 x $10^{-6}$ | $mm^2$ |
| Porosity | $\varphi$ | 0.7 | - |
| Capillary pressure (IPA 50%) | $P_{cap}$ | 3300 | Pa |
| Capillary pressure (nonanol) | $P_{cap}$ | 55004100 | Pa |
| Capillary pressure (pentanol) | $P_{cap}$ | 4500 | Pa |

[a] porosity and permeability are obtained from the literature[3], capillary pressures were determined by fitting $K$ and $\varphi$ to the experimental data.

### Section 1. Dynamics of capillary flow in the open-channel capillary tree

*Root channel.* We derive first the dynamics in the root channel. Neglecting the evanescent initial inertial regime,[9,10] the balance between capillary force $F_{cap}$ and wall friction $F_{drag}$ leads to

$$F_{cap} = p\,\gamma\,cos\theta^* = F_{drag} = p\,z_0\bar{\tau} = p\,z_0\left(\mu\frac{V_0}{\lambda}\right), \quad \text{(S1)}$$

where $\bar{\tau}$ is the average friction, $p$ the total channel perimeter in a cross section, $V_0$ the average velocity (which is a function of time and/or travel distance), λ the average friction length, $\mu$ the viscosity, $\gamma$ the surface tension, and $\theta^*$ the generalized Cassie angle[11] (in order to take into account the free surface of the open channel and accommodate for the potential of non-monolithic channels comprising different materials on the floor and walls). Then we have the relation between travel distance and time

$$\frac{d\,z_0^2}{dt} = \frac{2\,\lambda\,\gamma\,cos\theta^*}{\mu}, \tag{S2}$$

And finally

$$z_0 = \sqrt{\frac{2\,\lambda\,\gamma\,cos\theta^*}{\mu}}\,\sqrt{t} \tag{S3}$$

Note that the time at which the flow reaches the bifurcation at a distance $L_0$ from entrance, is

$$t_0 = \frac{\mu}{2\lambda\,\gamma\,cos\theta^*}\,L_0^2 = \frac{L_0^2}{C} \tag{S4}$$

where $C = \frac{2\,\lambda\,\gamma\,cos\theta^*}{\mu}$.

*Tree branches.* After the first bifurcation, at the level $n$, we must use a formulation that uses the pressures and write the pressure equilibrium along a fluidic path[1,12–15]

$$p_0\,L_0\,\mu\frac{V_0}{\lambda_0}\frac{1}{S_0} + \cdots + p_n\,z_n\,\mu\frac{V_n}{\lambda_n}\frac{1}{S_n} = \frac{p_n\,\gamma\,cos\theta_n^*}{S_n}, \tag{S5}$$

where $z_n$ here is the travel distance in the $n^{th}$ ramification, $S_n$ the cross-sectional area and $V_n$ the velocity in the daughter branch ($V_n = \frac{dz_n}{dt}$). Relation (S5) indicates that the capillary pressure is equal to the pressure associated to the friction along a path starting from the root channel to the $n^{th}$ channel. Using the relations linked to the homothetic ratio α, we have for the cross-sectional perimeters are

$$p_n = \alpha p_{n-1} = \cdots = \alpha^{n-1} p_1 = \alpha^n p_0, \tag{S6}$$

and the cross-sectional areas are

$$S_n = \alpha^2 S_{n-1} = \cdots = \alpha^{2(n-1)} S_1 = \alpha^{2n} S_0 \tag{S7}$$

Moreover, the friction lengths are homothetic because they are proportional to the hydraulic diameter of the channel

$$\lambda_n = \alpha \lambda_{n-1} = \cdots = \alpha^{n-1} \lambda_1 = \alpha^n \lambda_0 \tag{S8}$$

Hence the ratio $\frac{p_i}{\lambda_i}$ is constant for all indices $i$ from *0* to *n*, equal to $\frac{p_0}{\lambda_0}$. Moreover, the Cassie angles are everywhere the same, i.e., $\theta^* = \theta_1^* = \cdots = \theta_n^*$. Substituting (S6), (S7) and (S8) in (S5) yields

$$\alpha^{2n} L_0\, V_0 + \alpha^{2(n-1)}\, L_1\, V_1 + .. + \alpha^2 L_{n-1} V_{n-1} + \ z_n\, V_n = \frac{p_n}{p_0}\frac{\gamma}{\mu} \lambda_0\, cos\theta^* \ = \alpha^n \frac{\gamma}{\mu} \lambda_0\, cos\theta^* = \alpha^n \frac{C}{2}\,. \tag{S9}$$

Using the mass conservation equation,

$$V_0 = (2\,\alpha^2) V_1 = (2\,\alpha^2)^2\, V_2 \ = \cdots = (2\,\alpha^2)^n\, V_n, \tag{S10}$$

and remarking that $V_n = \frac{dz_n}{dt}$, (S9) becomes a differential equation in $z_n$ that can be integrated to obtain a quadratic polynomial in $z_n$

$$z_n^2 + 2[(2\alpha^4)^n L_0 + (2\,\alpha^4)^{n-1} L_1 + \cdots + 2\alpha^4 L_{n-1}] z_n = \alpha^n C\,(t - t_{n-1})\,, \tag{S11}$$

Denoting $A_n = (2\alpha^4)^n L_0 + (2\,\alpha^4)^{n-1} L_1 + \cdots + 2\alpha^4 L_{n-1} = (2\alpha^4)^n L_0 \Sigma_{n-1}$ , where $\Sigma_{n-1} = \left[1 + \frac{L_1}{2\alpha^4 L_0} + \cdots + \frac{L_{n-1}}{(2\,\alpha^4)^{n-1} L_0}\right]$, the solution is

$$z_n = A_n \left[-1 + \sqrt{1 + \frac{\alpha^n C}{A_n^2}\,(t - t_{n-1})}\right] \tag{S12}$$

Note that the travel distance given by relation (S12) is not of the conventional Lucas-Washburn form ($z \approx \sqrt{t}$), but of the form $z \approx -A + \sqrt{A^2 + \alpha^n C t}$ .

### Section 2. Dynamics of capillary flow in the paper pads (coupled to that in the open-channel capillary tree)

When reaching the extremities of the tree branches, the liquid start wicking the paper pads. The motion wicking of the pads is governed by Darcy's law [16,17]

$$V_p = -\frac{K}{\mu\phi}\,\nabla P = \frac{K}{\mu\phi}\,\frac{P_{cap} - P_j}{z_p}. \tag{S13}$$

where $P_{cap}$ is the capillary pressure of the paper and $P_j$ the pressure at the junction channel-paper pad. $K$ is the permeability of the pad and $\varphi$ its porosity. The index $p$ refers to the paper. The triplet ($P_{cap}$, $K$, $\varphi$) characterizes the paper strip.[18,19]

Three assumptions are used in the present model. First, the paper pads are homogeneous, i.e., there are no regions of higher or lower porosity. Hence, degree of saturation (local percentage of liquid) is assumed the same everywhere in the paper pad.[20,21] As a consequence, the capillary pressure $P_{cap}$ is constant everywhere, and there is no smearing of the advancing contact line, which is described in the literature as a sharp front approach.[22] Second, it is assumed that the dilatation of the paper fibers with the penetrating the liquid is negligible, so that the porosity $\varphi$ is constant everywhere in the pad. Third, the cellulose fibers do not absorb the wicking liquid, so that the mass conservation is independent of the time. Then equation (S13) can be cast under the form

$$\frac{d(z_p)^2}{dt} = \frac{2K}{\mu\phi}\left(P_{cap} - P_j\right). \qquad \text{(S14)}$$

In order to solve equation (S14), the pressure $P_j$ mut be determined. According to (S5)

$$P_j = p_0\, L_0\, \mu \frac{V_0}{\lambda_0}\frac{1}{S_0} + \cdots + p_n\, L_n\, \mu \frac{V_n}{\lambda_n}\frac{1}{S_n}\,, \qquad \text{(S15)}$$

where $n$ is the channel level at the junction with the paper pads. Using again (S6), (S7), (S8) and (S10), the pressure $P_j$ can be expressed as

$$P_j = p_0\, L_0\, \mu \frac{V_0}{\lambda_0}\frac{1}{S_0}\left(1 + \frac{L_1}{L_0}\frac{1}{2\,\alpha^4} + \cdots + \frac{L_n}{L_0}\frac{1}{(2\,\alpha^4)^n}\right) = p_0\, L_0\, \mu \frac{V_0}{\lambda_0}\frac{1}{S_0}\Sigma_n\ . \qquad \text{(S16)}$$

Using the mass conservation equation, the velocity $V_0$ can be expressed in function of the velocity in the paper

$$V_0 S_0 = 2^n \phi V_p S_p\ , \qquad \text{(S17)}$$

where $S_p$ is the total cross-sectional area of the pad. We consider two types of paper pads: rectangular and conical (triangular). We can group together the two geometries by writing

$$S_p = S_{p,0} + 2\beta\, h_p\, z_p\ , \tag{S18}$$

where $z_p$ is the penetration distance, $h_p$ is the thickness of the paper pad, $S_{p,0}$ the cross section of the paper at the junction—there can be a sharp increase of section between the last channels of the tree and the pads— and β the cone semi-angle. In the case of a rectangular pad, $\beta = 0$. In the case of a cone, the wetted cross-section in the paper pad is assumed to have a perfectly rounded interface.

Successively substituting (S18) in (S17) and (S17) in (S16) yields the expression of the pressure $P_j$ in function of $V_p$ and $z_p$

$$P_j \ = \frac{p_0\, L_0}{\lambda_0 S_0}\, \mu\, \Sigma_n\, 2^n \phi V_p \left(\frac{S_{p,0}}{S_0} + \frac{2\beta\, h_p\, z_p}{S_0}\right). \tag{S19}$$

For simplifying the notations, let us note $a_n = 2^n \Sigma_n K\, \frac{p_0 L_0}{\lambda_0\ S_0}$, $\delta = \frac{S_0}{2\,\beta\, h_p}$ and $b = \frac{2K}{\mu\phi} P_{cap}$. The two first parameters $a_n$ and $\delta$ have the dimension of a length, while the unit for $b$ is $mm^2/s$. Substituting (S19) in (S14), and using the relation $V_p = \frac{dz_p}{dt}$, produce the differential equation for the penetration distance $z_p$

$$\frac{d(z_p)^2}{dt}\left(1 + \frac{a_n}{\delta}\right) + 2a_n\, \frac{S_{p,0}}{S_0}\, \frac{dz_p}{dt} - b = 0. \tag{S20}$$

The solution is

$$z_p = \frac{a_n}{\left(1+\frac{a_n}{\delta}\right)} \frac{S_{p,0}}{S_0} \left( -1 + \sqrt{1 + \left(\frac{S_0}{S_{p,0}}\right)^2 \frac{b\left(1+\frac{a_n}{\delta}\right)\tau}{a_n^2}} \right) , \qquad \text{(S21)}$$

where $\tau = t - t_n$ is the time taken from the entrance of the pad ($\tau = 0$, $z_p = 0$). Note that, for rectangular pads, $\delta = \infty$, ($\frac{1}{\delta} = 0$), and (S21) becomes

$$z_p = a_n \frac{S_{p,0}}{S_0} \left( -1 + \sqrt{1 + \left(\frac{S_0}{S_{p,0}}\right)^2 \frac{b\,\tau}{a_n^2}} \right) . \qquad \text{(S22)}$$

For rectangular pads, it is verified that, if the capillary pressure $P_{cap}$ in the paper is very high (the parameter $b$ is large), (S22) can be simplified, and the travel distance in the paper is $z_p \approx \sqrt{bt} = \sqrt{\frac{2K\,P_{cap}}{\mu\phi} t}$, which is simply the Lucas-Washburn law for paper alone, obtained by direct integration of (S14) with $P_j = 0$. In this case, the influence of the tree completely disappears.

In the general case, deriving (S21) in respect to time, the velocity of the fluid in the pad is

$$V_p = \frac{S_0}{S_{p,0}} \frac{b}{2a_n \sqrt{1+\left(\frac{S_0}{S_{p,0}}\right)^2 \frac{b\left(1+\frac{a_n}{\delta}\right)\tau}{a_n^2}}} . \qquad \text{(S23)}$$

Using the mass conservation equation (S17), the velocity in the root channel when the liquid wicks the paper is given by

$$V_{root} = 2^n V_p \phi \frac{S_p}{S_0} = \phi\left(\frac{S_{p,0}}{S_0} + \frac{z_p}{\delta}\right)\frac{S_0}{S_{p,0}} \frac{b}{2a_n\sqrt{1+\left(\frac{S_0}{S_{p,0}}\right)^2 \frac{b\left(1+\frac{a_n}{\delta}\right)\tau}{a_n^2}}} , \quad \text{(S24)}$$

where $z_p$ is given by (S21). At the precise time when the fluid contacts the pad ($\tau = 0$, $z_p = 0$), relation (S24) yields $V_{root,0} = \frac{\phi\, b}{2a_n}$, and replacing $a_n$ and $b$ by their expressions, $\frac{P_{cap}}{\Sigma_n} = \frac{p_0 L_0}{S_0}\frac{\mu\, V_{root,0}}{\lambda_0}$, which is the expression of the pressure $P_0$ at the end of the root channel when the velocity is $V_{root,0}$.

Let us consider the case of conical pads. The penetration distance $z_p$ can be removed from (S24) by using (S21)

$$V_{root} = \frac{2^{n-1}\phi\, b}{a_n}\frac{\left(\delta + a_n\sqrt{1+D\tau}\right)}{(\delta + a_n)\sqrt{1+D\tau}} , \quad \text{(S25)}$$

where $D = \left(\frac{S_0}{S_{p,0}}\right)^2 \frac{b\left(1+\frac{a_n}{\delta}\right)}{a_n^2}$ .

The unit of $D$ is 1/s. In the case of rectangular pads, the derivation of $V_{root}$ yields

$$V_{root} = \frac{2^{n-1}\phi\, b}{a_n}\frac{1}{\sqrt{1+D_r\tau}} , \quad \text{(S27)}$$

where $D_r = \left(\frac{S_0}{S_{p,0}}\right)^2 \frac{b}{a_n^2}$

TABLE SIV. Summary of coefficients used in the different solvents

| Coefficient | Unit | IPA 50% | Nonanol | Pentanol |
|---|---|---|---|---|
| $a$ | mm | 0.0026 | 0.0026 | 0.0026 |
| $d$ | mm | 1.6 | 1.6 | 1.6 |
| $a/d$ | - | 0.002 | 0.002 | 0.002 |
| $Sp_0/Sp$ | - | 0.38 | 0.38 | 0.38 |
| $b$ | $mm^2/s$ | 3.7 | 1.8 | 2.0 |
| Viscosity $\mu$ | mPa.s | 3.2 | 11.2 | 3.75 |
| Capillary pressure $P_{cap}$ | Pa | 2072 | 4116 | 4500 |
| $2^n S_n$ | - | 11.34 | 11.34 | 11.34 |
| Surface tension $\gamma$ | mN/m | 24 | 28.5 | 25.5 |
| Contact angle $\theta$ | degrees | 18 | 13 | 10 |
| Cassie contact angle $\theta^*$ | degrees | 55.5 | 52.0 | 51.1 |
| Capillary force $\gamma \cos \theta^*$ | mN/m | 13.6 | 17.5 | 15.6 |
| Washburn coefficient $\sqrt{\left(\frac{\gamma}{\mu}\right) 2\lambda \cos\theta^*}$ | $mm/s^{1/2}$ | 47.0 | 28.5 | 38.7 |
| Washburn coefficient from fit with experiments $\sqrt{\left(\frac{\gamma}{\mu}\right) 2\lambda \cos\theta^*}$ | $mm/s^{1/2}$ | 38 | 25 | 40 |

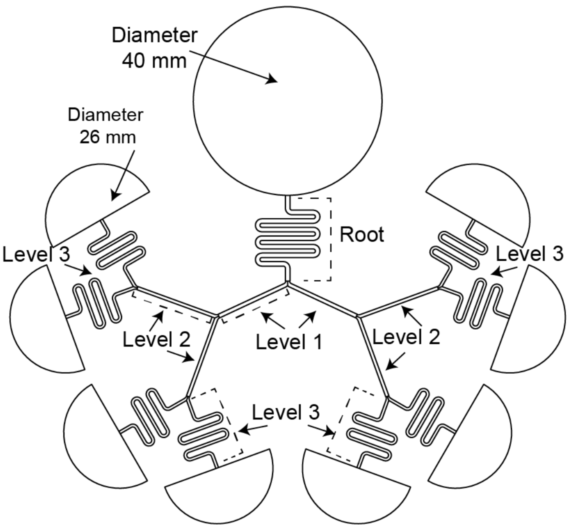


| Level (L) | Length (mm) | Width (mm) | Depth (mm) |
|---|---|---|---|
| 0 (root) | 98.0 | 1.06 | 1.76 |
| 1 | 18.2 | 0.90 | 1.50 |
| 2 | 20.1 | 0.77 | 1.28 |
| 3 | 58.8 | 0.65 | 1.08 |

FIG. SI. Engineering drawing of device with dimensions.

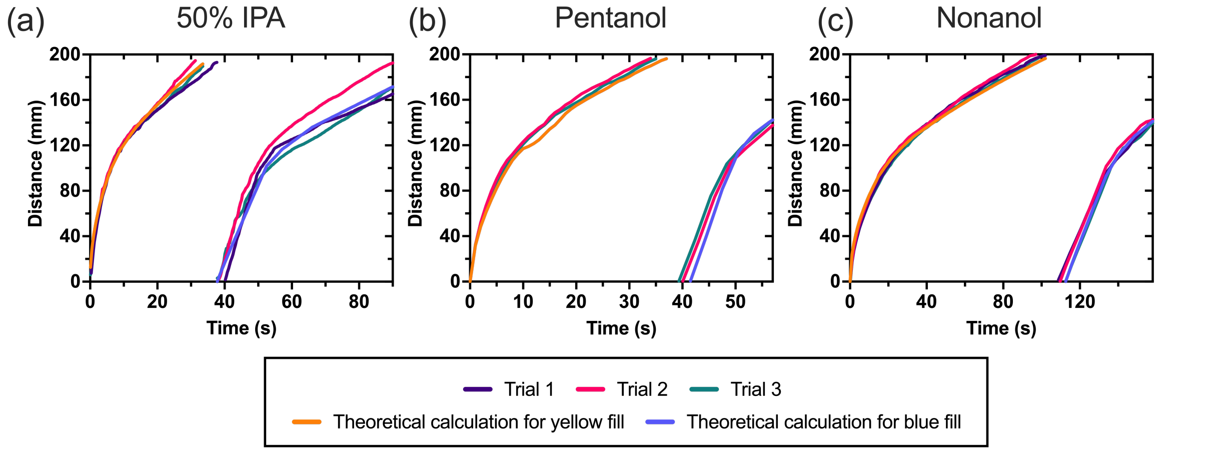


FIG. SII. Raw data plots of all trials for (a) 50% IP, (b) pentanol, and (c) nonanol of distance traveled over time.

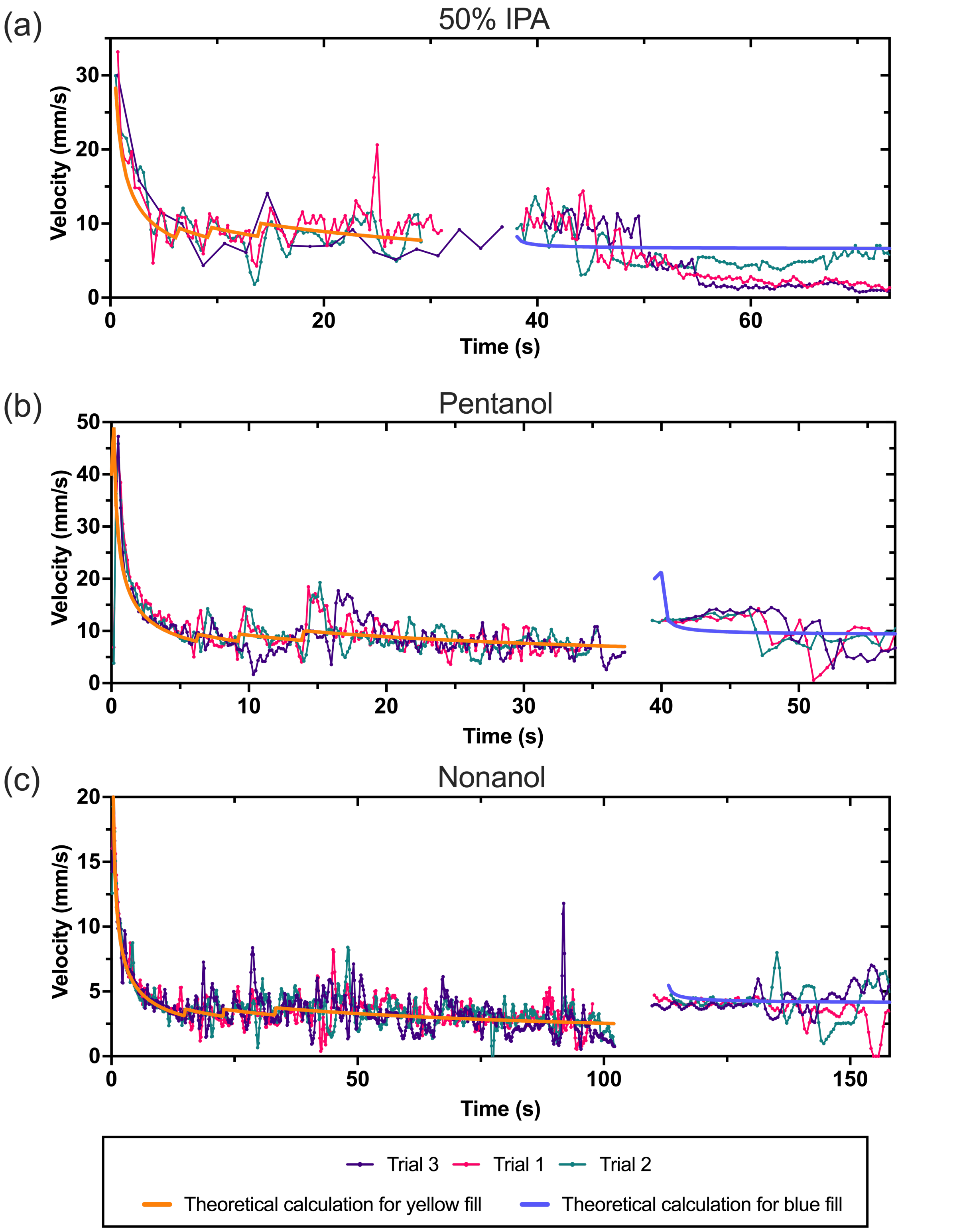


FIG. SIII. Raw data plot of all trials for (a) 50% IPA, (b) pentanol, and (c) nonanol of velocity over time.

†Authors contributed equally

[1] J. J. Lee, J. Berthier, A. B. Theberge, and E. Berthier, Langmuir 35, 10667 (2019).

[2] J. J. Lee, J. Berthier, K. E. Kearney, E. Berthier, and A. B. Theberge, Langmuir 36, 12795 (2020).

[3] M. Boodaghi and A. Shamloo, AIChE J. 66, e16756 (2020).

[4] A. A. Mohammad, K. H. A. E. Alkhaldi, M. S. AlTuwaim, and A. S. Al-Jimaz, J. Chem. Thermodyn. 74, 7 (2014).

[5] S. Kim, P. A. Thiessen, E. E. Bolton, J. Chen, G. Fu, A. Gindulyte, L. Han, J. He, S. He, B. A. Shoemaker, J. Wang, B. Yu, J. Zhang, and S. H. Bryant, Nucleic Acids Res. 44, (2016).

[16] H. Darcy, Les Fontaines Publiques de La Ville de Dijon, (Paris, 1856).
